# Investigation of microbial community interactions between lake Washington methanotrophs using genome-scale metabolic modeling

**DOI:** 10.1101/2020.02.21.958074

**Authors:** Mohammad Mazharul Islam, Tony Le, Shardhat Reddy Daggumati, Rajib Saha

**Author notes:** Correspondence: Rajib Saha, Assistant Professor, Chemical and Biomolecular Engineering, University of Nebraska-Lincoln, Lincoln, NE-68588, USA. These authors contributed equally to this work (Tony Le, Shardhat Reddy Daggumati).

## Abstract

**Background:** The role of methane in global warming has become paramount to the environment and the human society, especially in the past few decades. Methane cycling microbial communities play an important role in the global methane cycle, which is why the characterization of these communities is critical to understand and manipulate their behavior. Methanotrophs are a major player in these communities and are able to oxidize methane as their primary carbon source.

**Results:** Lake Washington is a freshwater lake characterized by a methane-oxygen countergradient that contains a methane cycling microbial community. The major microbial members include methanotrophs such as *Methylobacter Tundripaludum 21/22* and *Methylomonas sp. LW13*. In this work, these methanotrophs are studied via developing highly curated genome-scale metabolic models. Each model was then integrated to form a community model with a multi-level optimization framework. The metabolic interactions for the community were also characterized. While both organisms are competitors for methane, *Methylobacter* was found to display altruistic behavior in consuming formaldehyde produced by *Methylomonas* that inhibits its growth. The community was next tested under carbon, oxygen, and nitrogen limited conditions to observe the systematic shifts in the internal metabolic pathways and extracellular metabolite exchanges. Each condition showed variable differences within the methane oxidation pathway, serine cycle, pyruvate metabolism, and the TCA cycle as well as the excretion of formaldehyde and carbon di-oxide from the community. Finally, the community model was simulated under fixed ratios of these two members to reflect the opposing behavior of the community in synthetic and natural communities. The simulated community demonstrated a noticeable switch in intracellular carbon metabolism and formaldehyde transfer between community members in natural vs. synthetic condition.

**Conclusion:** In this work, we attempted to reveal the response of a simplified methane recycling microbial community from Lake Washington to varying environments and also provide an insight into the difference of dynamics in natural community and synthetic co-cultures. Overall, this study lays the ground for *in silico* systems-level studies of freshwater lake ecosystems, which can drive future efforts of understanding, engineering, and modifying these communities for dealing with global warming issues.

## Introduction

The accelerated rise in worldwide average temperature in recent years is posing a serious threat to the environment, terrestrial ecosystems, human health, economy, and the ultimate survival of the planet earth. About 20 percent of global warming is caused by methane and it is expected to be 86 times more potent than carbon di-oxide in warming the earth over the next two decades (Houghton, Jenkins & Ephraums, 1990; IPCC, 2013). The impacts of the rapid increase in atmospheric methane (Nisbet et al., 2019) are compounded as higher temperatures are associated with an increase in methane production from wetlands and lakes (Yvon-Durocher et al., 2014). Methanotrophs are gram-negative Proteobacteria that are an integral part of the global carbon cycle (Hanson & Hanson, 1996). They exist in diverse environments such as wetlands, lakes, and the tundra and use the enzyme methane monooxygenase (MMO) to oxidize methane as their sole source of carbon (Hanson & Hanson, 1996). Thus, methanotrophs act as the primary biological sink for methane (Hanson & Hanson, 1996), consuming up to 90 percent of the methane produced in soil/sediments in addition to the atmospheric methane (Whalen & Reeburgh, 1990; Krause et al., 2017). Methanotrophs have also shown the ability to produce various useful products such as single-cell proteins, biodiesels, biopolymers, and osmo-protectants (Strong, Xie & Clarke, 2015).

In 1970 over a hundred species of methanotrophs were isolated and their physical and physiological properties were characterized (Whittenbury & Wilkinson, 1970). Over time, the taxonomy of methanotrophs were revised utilizing physiological tests, morphological observations, numerical taxonomy, DNA-DNA hybridization analysis, and PLFA profiles (Bowman et al., 1993) Based on the metabolic pathways utilized to assimilate formaldehyde produced from oxidizing methane, methanotrophs are categorized into three major types. Type I methanotrophs assimilate formaldehyde using the ribulose monophosphate pathway, Type II methanotrophs assimilate formaldehyde using the serine cycle while Type X methanotrophs can use both (Hanson & Hanson, 1996).

Lakes act as a major source and sink for methane and account for 6 to 16 percent of biologically produced methane (IPCC, 2013; Yvon-Durocher et al., 2014). Lake Washington is a freshwater lake characterized by a methane-cycling community where various methanotrophs are the major microbes (Yu et al., 2016). It contains a steep vertical counter-gradient of methane and oxygen, and is separated into oxic and anoxic layers where methane production and consumption occur, respectively (Yu et al., 2016). Hence, it can be a model system to better understand methane-cycling communities in lakes and their role in the global methane cycle. Understanding the metabolic interactions in these communities will aid in developing methods to reduce the amount of methane emitted from lakes. A diverse array of microbes exist in the Lake Washington community, where Proteobacteria comprise 33% of the community and includes a major subtype of type I methanotrophs, *Methylococcaceae* at a 10% abundance level (Beck et al., 2013). The genus *Methylobacter* is the most dominant player in *Methylococcaceae* group at 47.7%, while other major players include *Crenothrix, Methylomonas*, and *Methylomicrobium* at 30.0%, 10.8%, and 7.4%, respectively (Beck et al., 2013). In addition, the genus *Methylocystaceae* represents the type II methanotrophs in the community, which are not highly abundant (Beck et al., 2013). Other members include organisms containing chloroplast sequences, bacteriodetes, acidobacteria, and chloroflexi (Beck et al., 2013).

Understanding the physiological dynamics and interactions in natural methane-cycling communities, such as the Lake Washington, are crucial to addressing problems concerning methane’s role in global warming and leveraging methanotrophs’ possible functions in bioremediation and bioproduction. Omics-based techniques and high-throughput sequencing can elucidate important features of the community such as taxonomic information, community composition, and presence of functionally important genes (Temperton & Giovannoni, 2012). However, it is difficult to assign functionality to members of the community and decipher the roles of individual players due to the complexity of the community and the data involved (Zengler, 2009; Zengler & Palsson, 2012). On the other hand, synthetic communities were proven to be efficient models to provide insight into metabolic capabilities and interactions (De Roy et al., 2014). Simple representative community structures can be made to lower complexity, achieve more consistent results, and efficiently elucidate inter-species interactions (De Roy et al., 2014). Certain Lake Washington community members are easier to cultivate in a laboratory setting than other members. For instance, *Methylomonas* and *Methylosinus* species previously shown ease of cultivation in the laboratory (Auman et al., 2000). However, *Methylobacter* species were difficult to isolate and demonstrated poor growth compared to *Methylomonas* and *Methylosinus* (Yu et al., 2016). A synthetic community comprising 50 different Lake Washington microbes belonging to 10 methanotrophic, 36 methylotrophic, and 4 non-methanotrophic heterobacteria showed that *Methylobacter* was outperformed by other members in the community (Yu et al., 2016). This was inconsistent with the stable isotype probing study that found *Methylobacter* is the dominant species in Lake Washington (Beck et al., 2013). A simple three-member synthetic community of *Methylobacter, Methylomonas*, and *Methylosinus* showed similar results in which other members dominated *Methylobacter* (Yu et al., 2016). These inconsistencies indicate that the complexities of biological systems often make it challenging to understand the functions and interactions within and among organisms in synthetic communities via *in vitro* and *in vivo* studies.

*In silico* evaluation and analysis utilizing mathematical relation-based modeling allow for a high-resolution understanding of the biological processes in a microbial community. The availability of genome-scale metabolic network models combined with biological constraints provide multiple methods to analyze, perform *in silico* experiment, develop hypotheses, and redesign biological systems at a genome-level (Zomorrodi & Maranas, 2012; Zomorrodi, Islam & Maranas, 2014; Maranas & Zomorrodi, 2016; Islam & Saha, 2018). To develop effective multi-species community models, significant and comprehensive knowledge of inter-species interactions and experimental data must be utilized. Compartmentalized community metabolic models were used to model simple microbial consortia involved in bioremediation, synthetic auxotrophic co-growth, human gut microbiome, and soil bacterial ecosystems (Stolyar et al., 2007; Bizukojc et al., 2010; Lewis et al., 2010; Zhuang et al., 2011; Shoaie & Nielsen, 2014; Henry et al., 2016). On the other hand, community modeling frameworks incorporating the trade-offs between species- and community-level fitness successfully modeled steady-state and dynamic behavior in naturally occurring and synthetic soil microbial communities, synthetic co-cultures for bioproduction, human gut microbiome, and very recently to understand the microbial interactions in bovine rumen and viral influences (Zomorrodi & Maranas, 2012; Zomorrodi, Islam & Maranas, 2014; Chan, Simons & Maranas, 2017; Islam et al., 2019). Other novel methods were also proposed for modeling of such communities involving elementary mode analysis, evolutionary game theory, nonlinear dynamics, and stochastic processes (Vallino, 2003; Lehmann & Keller, 2006; Shou, Ram & Vilar, 2007; Borenstein & Feldman, 2009; Freilich et al., 2009; Frey, 2010; Nadell, Foster & Xavier, 2010; Magnúsdóttir, Heinken & Kutt, 2017).

In this work, we developed a simplified community metabolic model with two representative and functionally important strains of Lake Washington, namely, *Methylobacter Tundripaludum 21/22* (hereafter, *Methylobacter*) and *Methylomonas sp. LW13* (hereafter, *Methylomonas*) as representative organisms of the major methane-oxidizing microbes in the Lake Washington ecosystem. These species were chosen because of their high abundance in Lake Washington, the ability to mitigate common pollutants, and produce desirable biological products, and the availability of genome-annotation. Draft models of these species were reconstructed followed by careful curation to ensure proper representation of the species. Metabolic pathways and individual reactions that are fundamental to the growth of these organisms such as the ribulose monophosphate pathway, pentose phosphate pathway, methane metabolism, amino acid synthesis and utilization, serine cycle, and calomide biosynthesis were manually scrutinized and then integrated into the models. The interactions between the two models were established by referring to literature and known transported metabolites. The curated models of *Methylobacter* (704 genes, 1407 metabolites, and 1484 reactions) and *Methylomonas* (657 genes, 1454 metabolites, and 1464 reactions) were then utilized to develop a community model using a multi-level optimization framework, which was used to estimate biologically feasible metabolite secretion profiles and community compositions (Zomorrodi & Maranas, 2012; Zomorrodi, Islam & Maranas, 2014; Islam et al., 2019). The community was placed under carbon, oxygen, and nitrogen-limiting environments to study the changes in intracellular carbon and nitrogen metabolism and metabolite excretion profiles. The community composition of carbon-limited environments shows a shift in the carbon metabolism of both species as they scavenge carbon dioxide using components of the serine cycle. Similarly, the community exhibits commensal relationship in oxygen limiting condition, when formaldehyde is transferred to *Methylobacter* from *Methylomonas.* This helps *Methylobacter* to dominate the community and safeguard *Methylomonas* from the toxicity and growth inhibitory effects of formaldehyde (Hou, Laskin & Patel, 1979; Costa et al., 2001). Our results also indicate metabolic reprogramming in TCA cycle and pyruvate metabolism, which can help generate new hypotheses for *in vivo* experiments. We also simulated the observed binary compositions in natural community and synthetic co-culture to elucidate the changes in intra- and extracellular metabolic fluxes, specifically a shift to formaldehyde transfer between organisms instead of the primary carbon source methane. Overall, our results enhance the mechanistic understanding of the Lake Washington methane-cycling community, which can drive further engineering efforts for efficient rerouting of carbon and nitrogen as well as mitigation of methane emission from freshwater ecosystems globally.

## Materials & Methods

### Metabolic model reconstruction

The draft genome-scale metabolic models of *Methylobacter* and *Methylomonas* were developed and downloaded using the ModelSEED database (Henry et al., 2010). The models included reactions for glycolysis/gluconeogenesis, citrate cycle, pentose phosphate pathway, steroid biosynthesis, nucleotide metabolism, and various amino acid biosynthesis. Flux Balance Analysis (FBA), a mathematical approach for analyzing the flow of metabolites through a metabolic network, was utilized for model testing and analyzing flux distributions throughout the work (Orth, Thiele & Palsson, 2010). FBA implements the following optimization framework.

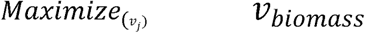

*subject to*

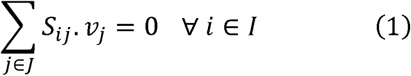

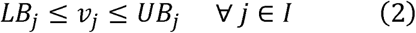

In the framework, *I* and *J* represent the sets of metabolites and reactions in the metabolic model, respectively. *S*_*ij*_ represents the stoichiometric coefficient of metabolite *i* in reaction *j*. The flux value of each reaction *j, v*_*j*_, must be within the parameters of the minimum, *LB*_*j*_, and maximum, *UB*_*j*_, biologically allowable fluxes. *v*_*biomass*_ is the flux of the biomass reaction which simulates the yield of cellular growth in the model (Orth, Thiele & Palsson, 2010).

### Metabolic Model Curation

Draft models for both *Methylobacter* and *Methylomonas* underwent an extensive manual curation process, including chemical and charge-balancing, elimination of thermodynamically infeasible cycles, and ensuring network connectivity. Reactions in the draft model reaction set imbalanced in protons were checked with their appropriate protonation states and corrected by adding and deleting proton(s) on either side of the reaction equation. The remaining imbalanced reactions were stoichiometrically inaccurate and required the atoms on both sides of the reaction equation to be balanced.

While applying mass balance constraints to genome-scale metabolic models can display the net accumulation and consumption of metabolites within each microbial model, it fails to account for the regulation of reaction fluxes. The limitation of this constraint is better elucidated when focusing on reaction cycles that do not consume and produce metabolites. Because of the absence of metabolite consumption and production, the overall thermodynamic driving force of the cycles become zero and the cycle is incapable of supporting any net flux, and thus deemed thermodynamically and biologically infeasible (Schellenberger, Lewis & Palsson, 2011). These thermodynamically infeasible cycles in our models were identified by inhibiting all nutrient uptakes to the cell and utilizing the optimization framework, Flux Variability Analysis (FVA), which maximizes and minimizes each reaction flux within the model based on the mass balance constraints (Mahadevan & Schilling, 2003). Reactions whose fluxes reached the defined lower and upper bounds were determined to be unbounded reactions, group together based on stoichiometry, and systematically corrected. These cycles were eliminated by removing duplicate reactions, turning off lumped reactions, fixing reaction directionality, or selectively turning reactions on or off based on cofactor specificities found from literature (Ishida et al., 1969; Dekker, Lane & Shapley, 1971; Chen, Lee & Chang, 1991; Schomburg & Stephan, 1995; Achterholt, Priefert & Steinbuchel, 1998; Hadfield et al., 1999; Hutter & Singh, 1999; Kai, Matsumura & Izui, 2003; Feist et al., 2007; Dean, 2012; Flamholz et al., 2012; Lin et al., 2014; Christensen et al., 2017). The FVA optimization algorithm is shown below.

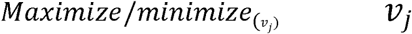

*subject to*

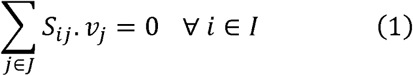

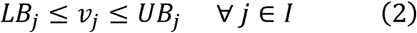

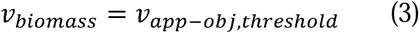

Both metabolic models were checked for the ability to produce biomass and metabolites they were known to produce (Svenning et al., 2011; Kalyuzhnaya et al., 2015). The metabolic functionalities of the models were ensured by identifying and manually adding reactions from biochemical databases, such as KEGG (Kanehisa & Goto, 2000) and Uniprot (Consortium, 2018), to each model. Fully developed models of related organisms such as *Methylococcus capsulatus* and *Methanomonas methanica* for both *Methylobacter* and *Methylomonas* were utilized in help pinpoint absent enzymatic activity within our models. The addition of these reactions was confirmed using the bioinformatic algorithm, BLAST, which compares the genes of our organisms and related organisms and determines whether they are found to be orthologous. It was then tested whether these reactions would increase the number of thermodynamically infeasible cycles and promote for further curation of the models.

### Community model formation

Following the curation of each individual microbial model, both were implemented to form a community model using the bi-level multi-objective optimization framework OptCom (Zomorrodi & Maranas, 2012). OptCom simultaneously optimizes each individual community member’s flux balance analysis problem as an inner-level optimization problem and the community model objective as an outer-level optimization problem. The mathematical framework of the OptCom procedure is the following.

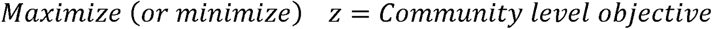

*subject to*,

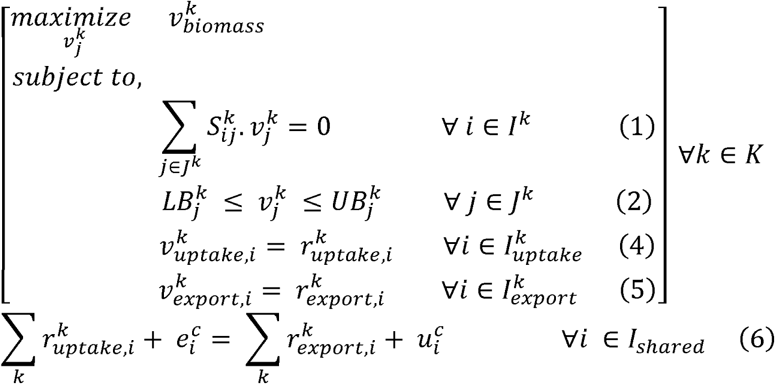

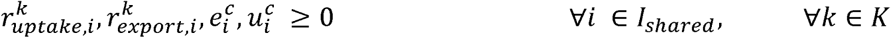

The inner-level optimization problem(s) represents the steady-state FBA problem for each community member *k* and limits the exchange (uptake and export) fluxes of shared metabolites between the individual species using the outer-level optimization problem parameters *r*^*k*^_*uptake,i*_ and *r*^*k*^_*export,i*_, respectively. The outer-level optimization problem constraint characterizes the mass balance for every shared metabolite in the extracellular environment within the shared metabolite pool. Metabolic interactions, with constraints, were confirmed with prior experimental research involving both community members or one member with a related organism found in the Lake Washington community (Yu et al., 2016). Incorporating these known interactions provides a stronger elucidation of how these organisms interact in their environment.

### Formaldehyde Inhibition Constraint

Formaldehyde production can be inhibitory towards *Methylomonas* growth (Bussmann, Rahalkar & Schink, 2006). *Methylobacter* is able to uptake formaldehyde secreted from *Methylomonas* to alleviate inhibitory effects on *Methylomonas* growth. A constraint based upon the minimum and maximum inhibitory concentrations of formaldehyde was implemented to simulate formaldehyde’s inhibitory effects on *Methylobacter* (Hou, Laskin & Patel, 1979; Costa et al., 2001). The minimum formaldehyde concentration required for growth inhibition was found to be 1 mM (Hou, Laskin & Patel, 1979) and the formaldehyde concentration required for total growth inhibition (maximum formaldehyde concentration) was found to be 7 mM (Costa et al., 2001).

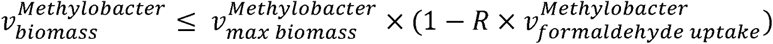

The parameter *R* was derived by finding the rate of change of biomass growth by the change in formaldehyde concentration.

### Identification of unknown community interactions and connecting metabolic network gaps

Automated draft reconstructions are limited as many reaction networks possess gaps due to missing reactions and blocked reactions. These are defined as reactions that lack production/consumption of its reactants/products. Major metabolic pathways were added based upon the literature of each organism (Kalyuzhnaya et al., 2015). Gaps were filled by referencing genetically related organisms to find missing metabolic capabilities. The presence of these possible reactions in the models were validated by cross referencing the relevant amino acid sequences between the reference organism and our models via pBlast. The reactions were are then checked for the formation of thermodynamically infeasible cycles before being accepted. Metabolic interactions between the community members were identified through existing literature (Hou, Laskin & Patel, 1979; Costa et al., 2001; Bussmann, Rahalkar & Schink, 2006; Kalyuzhnaya et al., 2015).

### Computational resources

The General Algebraic Modeling System (GAMS) version 24.8.5 with IBM CPLEX solver was utilized to run the optimization algorithms. The optimization frameworks were scripted in GAMS and then run on a Linux-based high-performance cluster computing system at the University of Nebraska-Lincoln. The downloaded models from ModelSEED with curations were parsed from System-Biology Markup Language (SBML) level 2 documentation using general-purpose programming language Python to generate the input files necessary for GAMS and subsequent manual curations.

## Results

### Genome-scale metabolic models of *Methylobacter* and *Methylomonas*

The draft genome-scale models of *Methylobacter and Methylomonas* are reconstructed using the ModelSEED database (Henry et al., 2016). The manual curation process ensures that there is no chemical and charge imbalance present in either of the models and there is no reaction with unrealistically high fluxes (infeasible reactions) without any nutrient uptake. The manual curation also reconnects a significant number of blocked metabolites to the network in both models (*i.e*., 99 metabolites for *Methylobacter* and 96 metabolites for *Methylomonas*). This enhancement of network connectivity is performed using available literature pertaining to major metabolic pathways that are known to be present in both the microbes (Kanehisa & Goto, 2000; Kalyuzhnaya et al., 2015). The draft models were lacking some reactions in the important metabolic pathways *i.e*., the methane oxidation, pentose phosphate pathway, nitrogen fixation, cofactor, and amino acid production, which are curated at this stage. The model statistics are shown in Table 1. Supplementary information 1 and 2 contain the model files for *Methylobacter* and *Methylomonas*, respectively.

**Table 1:** Model statistics for *Methylobacter* and *Methylomonas*.

| Models | *Methylobacter* | *Methylomonas* |
| --- | --- | --- |
| Genes | 704 | 657 |
| Reactions | 1484 | 1464 |
| Metabolites | 1407 | 1454 |
| Unbounded reactions | 0 | 0 |
| Blocked reactions | 688 | 687 |

### Community dynamics under variable environmental conditions

The individual models are integrated into a community model using available multi-objective computational optimization framework (Zomorrodi & Maranas, 2012). The metabolic interactions between *Methylobacter* and *Methylomonas* are described using inter-organisms flow constraints. The community, as a whole, consumes methane, oxygen, and nitrogen, which is then shared between *Methylobacter* and *Methylomonas*. In addition, *Methylobacter* consumes formaldehyde produced by *Methylomonas*, which alleviates the formaldehyde toxicity on *Methylomonas* growth (Bussmann, Rahalkar & Schink, 2006). At the same time, both *Methylobacter* and *Methylomonas* export carbon di-oxide to the environment. The community is simulated under four conditions in which methane, oxygen, and nitrogen are supplied to the community at various levels. These conditions are denoted as A, B, C, and D in Table 2 and correspond to the Figures 3,4, 5, 6, and 7, respectively. It should be noted that the amount of each nutrient consumed by the members are not necessarily equal to the amount of nutrient supplied as one of the nutrients acts as a limiting reagent in each limiting condition. The community model is illustrated in Figure 1.

**Table 2:** High and limiting nutrient conditions for community simulation.

|  |  |  |  |
| --- | --- | --- | --- |
| Condition | Methane uptake<br>(mmol/gDCW.hr) | Oxygen uptake<br>(mmol/gDCW.hr) | Nitrogen uptake<br>(mmol/gDCW.hr) |
| A (high C, high O, high N) | 100 | 100 | 100 |
| B (high C, limiting O, high N) | 100 | 10 | 100 |
| C (high C, high O, limiting N) | 100 | 100 | 1.2 |
| D (limiting C, high O, high N) | 16 | 100 | 100 |

**Figure 1.**
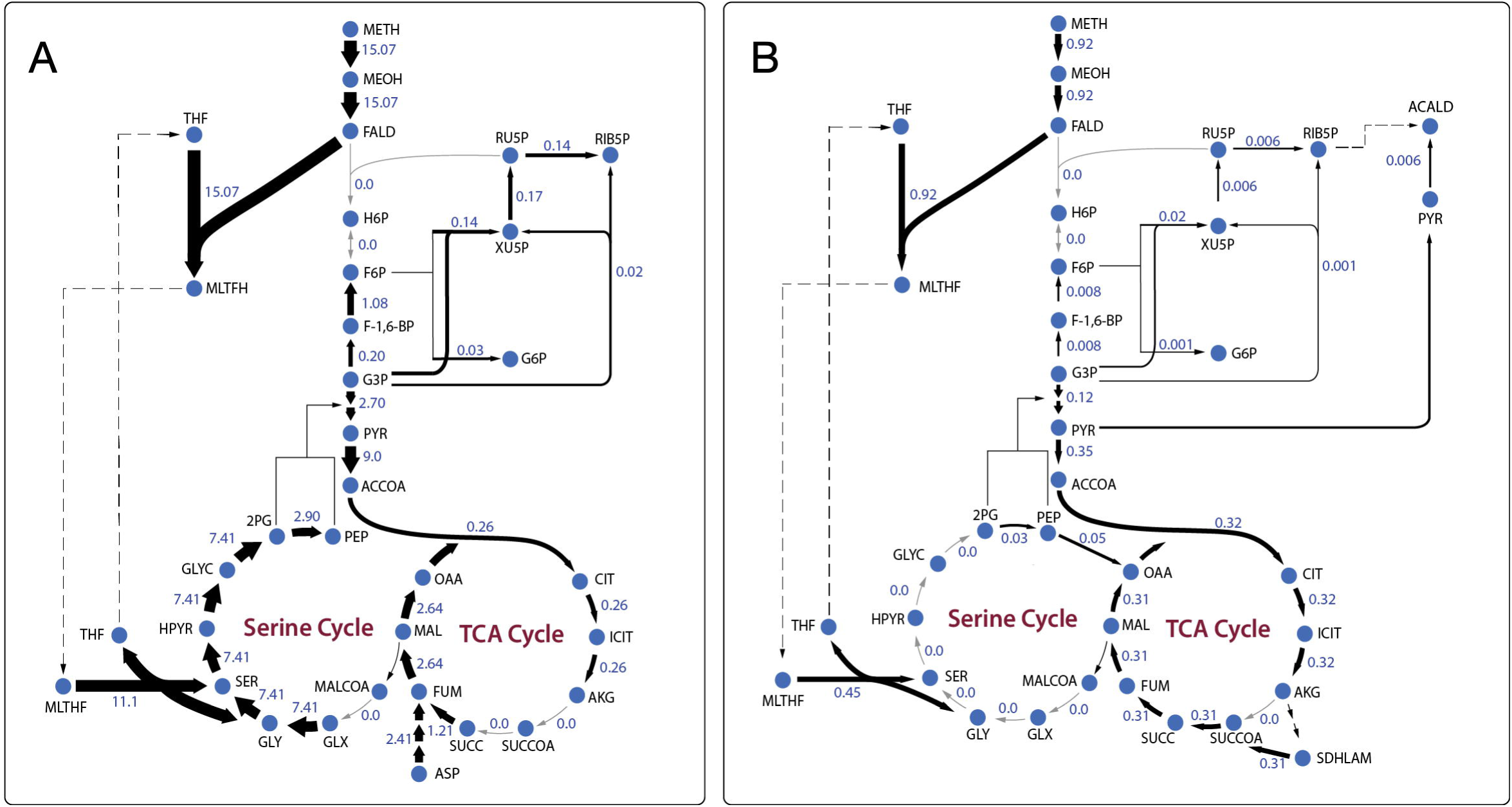
Community dynamics showing the fluxes of key shared metabolites and biomass in the community model. A) High nutrient condition; B) Oxygen limited condition; C) Nitrogen limited condition; and D) Carbon limited condition

**Figure 2:**
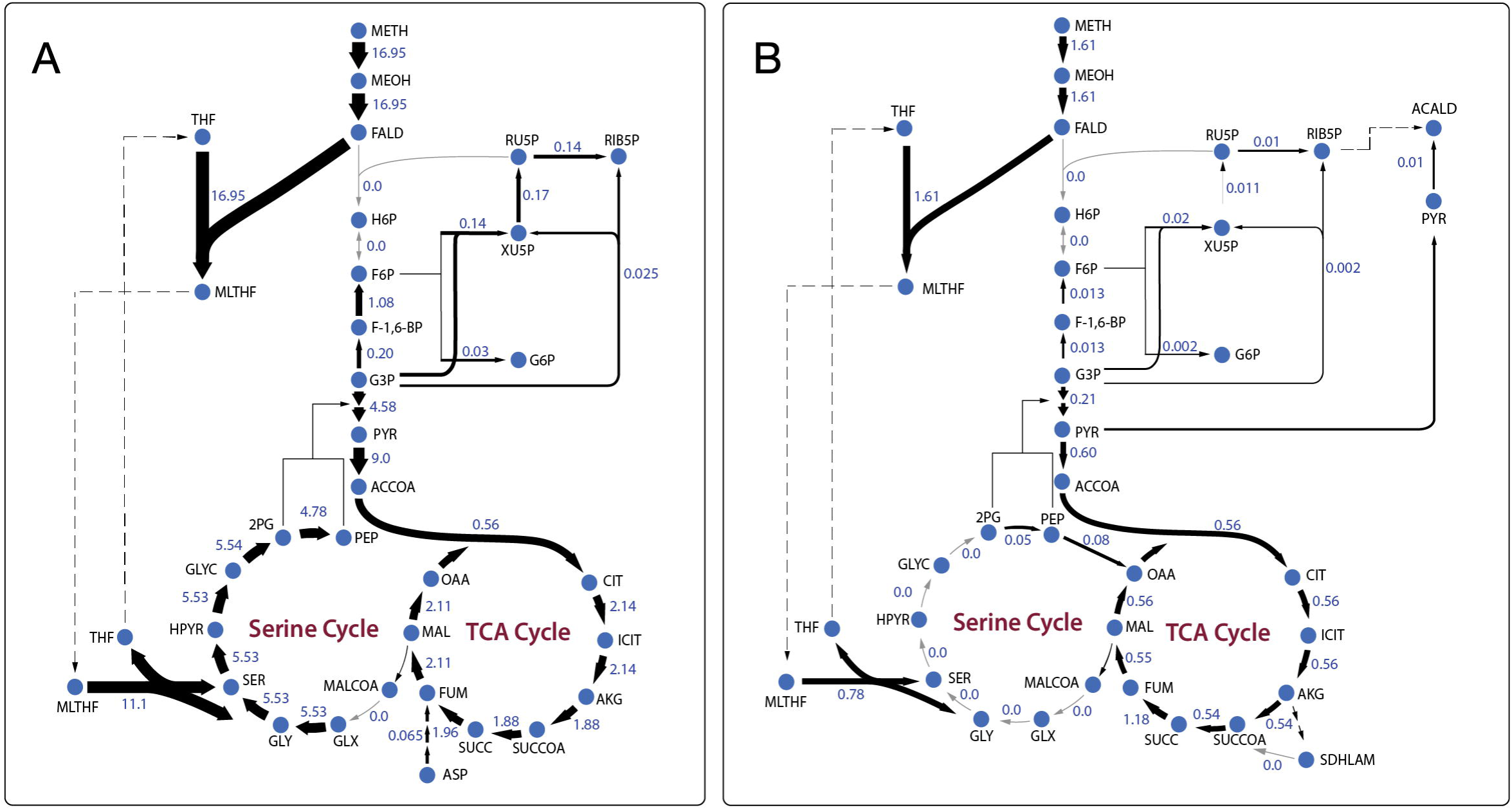
The community composition and total biomass under varying Methane and Oxygen conditions. The size of each pie chart represents the total community biomass.

**Figure 3:**
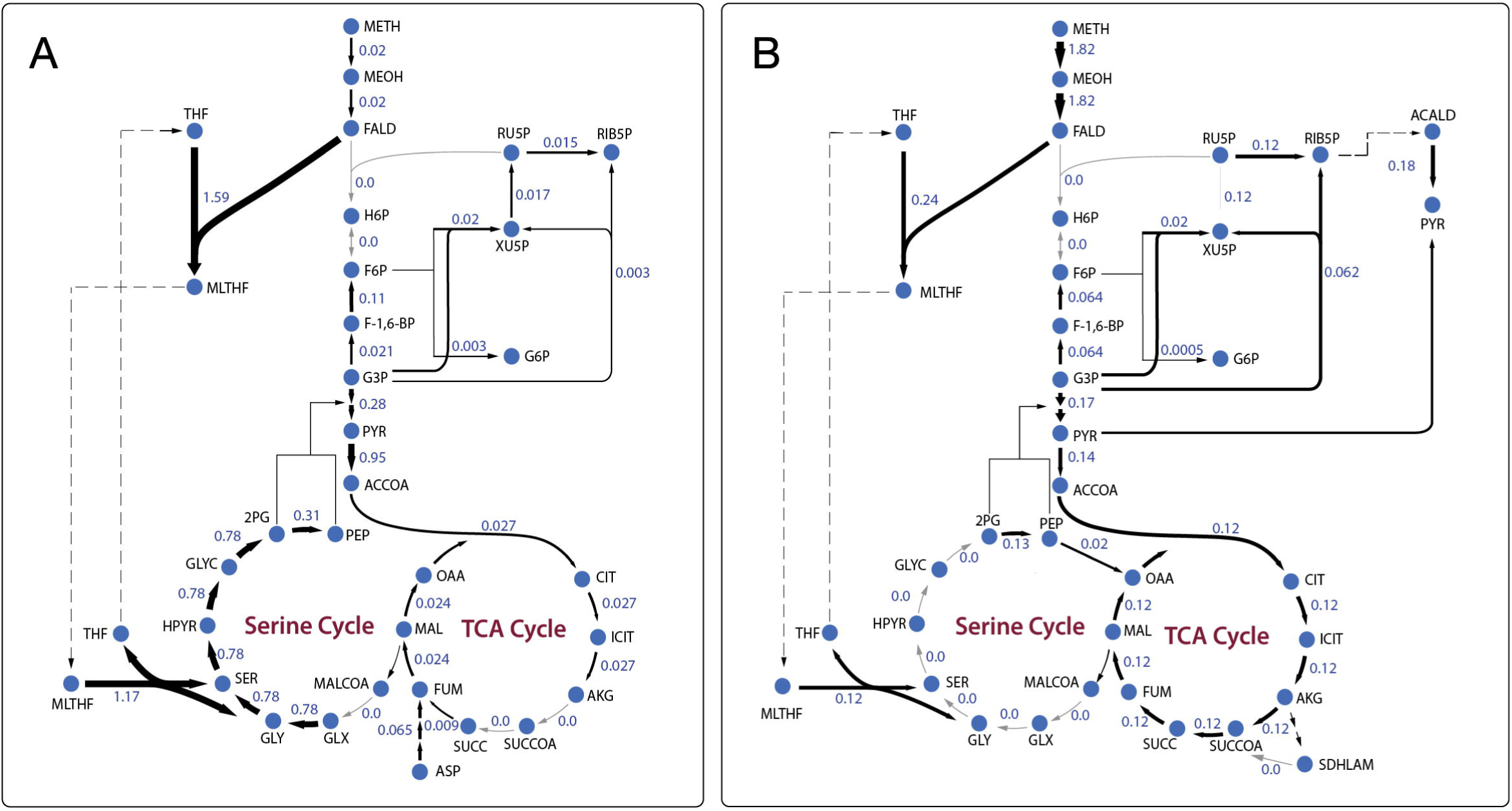
Central carbon metabolism fluxes in *Methylobacter* (A) and *Methylomonas* (B) under high nutrient condition.

**Figure 4:**
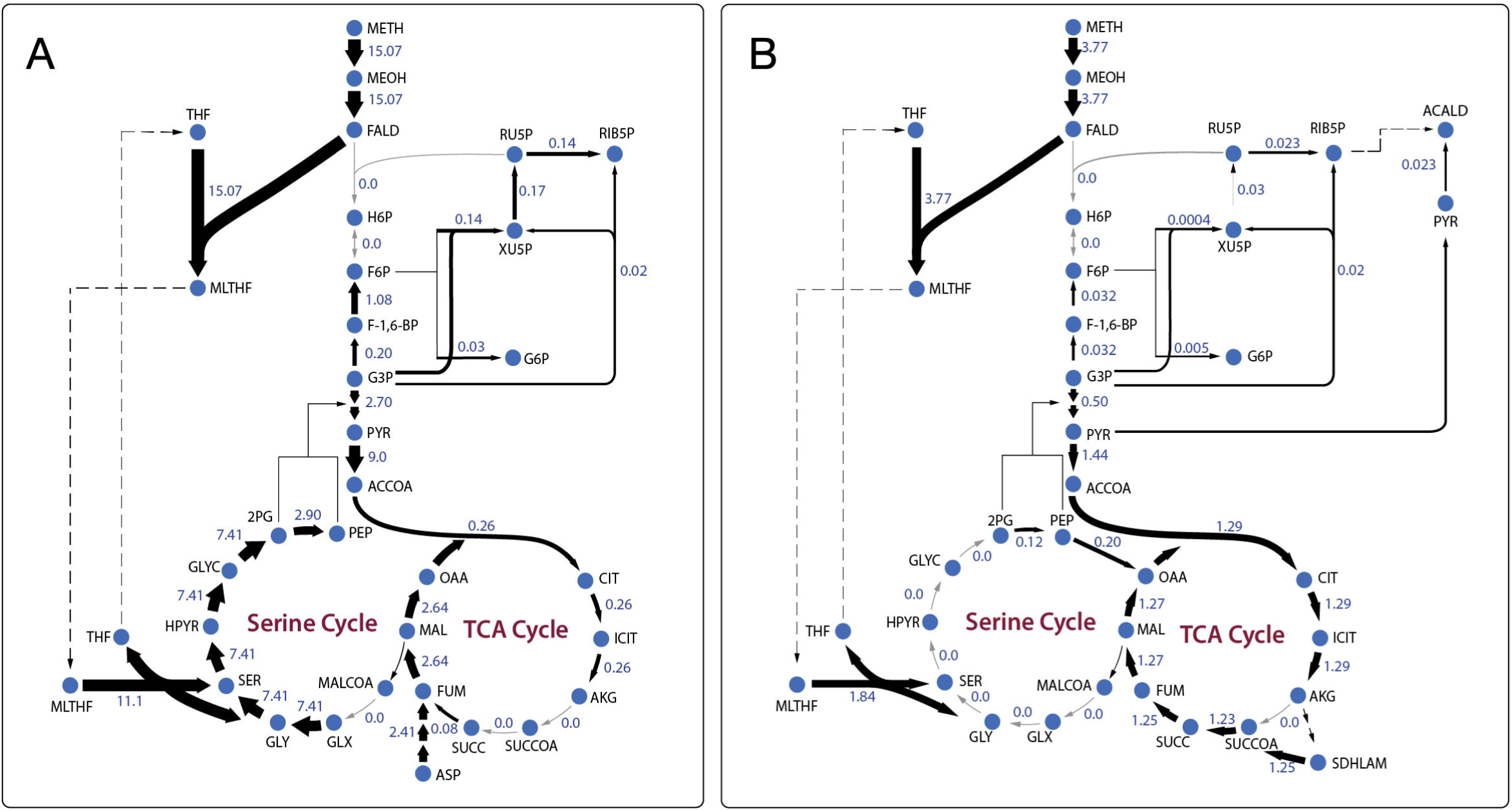
Central carbon metabolism fluxes in *Methylobacter* (A) and *Methylomonas* (B) under Oxygen limited condition.

**Figure 5:**
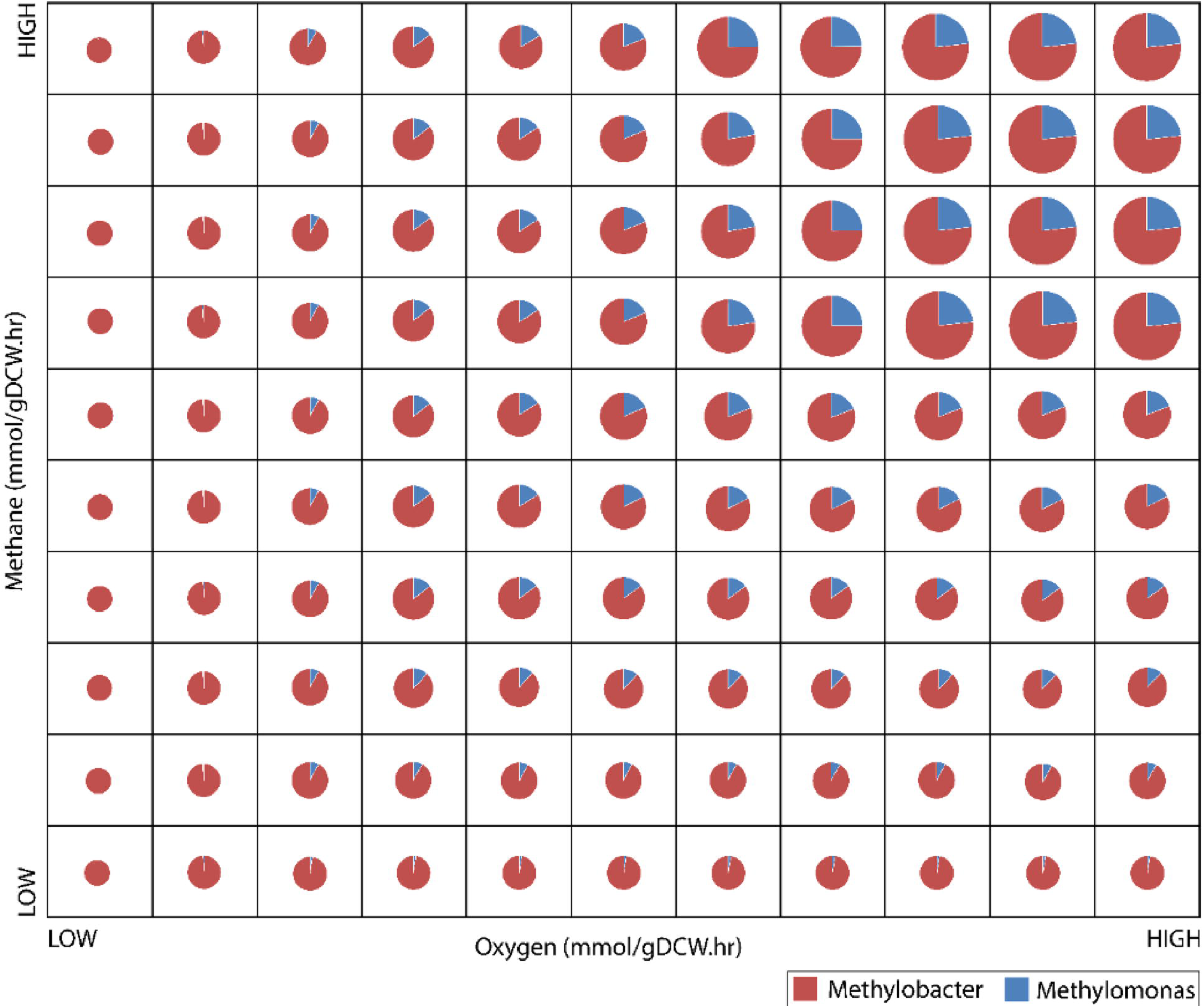
Central carbon metabolism fluxes in *Methylobacter* (A) and *Methylomonas* (B) under Nitrogen limited condition.

**Figure 6:**
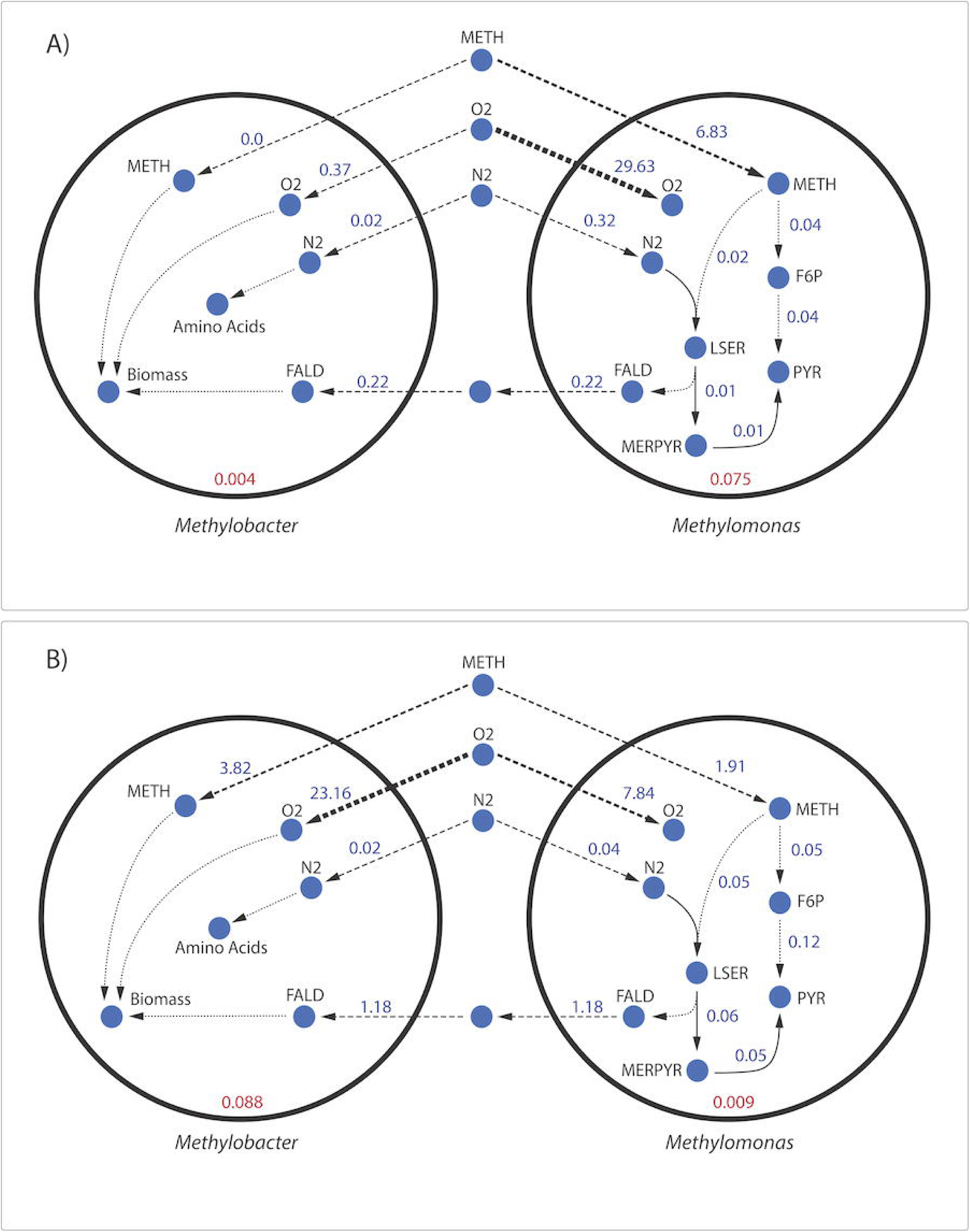
Central carbon metabolism fluxes in *Methylobacter* (A) and *Methylomonas* (B) under Carbon limited condition.

The community biomass flux is the highest when all the nutrients are highly abundant (Figure 1A). It is observed that a limited uptake of oxygen also imposes limits on the carbon and nitrogen utilization by the community, which results in reduced biomass fluxes for both *Methylomonas* and *Methylobacter* (Figure 1 B). When oxygen uptake is limited, methane utilization by *Methylomonas* is higher compared to *Methylobacter* despite an overall decrease in methane consumption by the community. Only in the oxygen limiting condition formaldehyde transfer from *Methylomonas* to *Methylobacter* is observed, which indicates that due to the oxygen limitation, carbon flux is shifted from biomass towards formaldehyde production. As expected, this methanotrophic community produces carbon dioxide as a byproduct of their methane oxidation at all conditions, with a lower rate in oxygen-limiting condition (Figure 1B). The results are presented in Supplementary information 3 in detail.

The total community biomass and community composition under methane and oxygen gradients are simulated to model the methane-oxygen counter gradient that exists in Lake Washington (Yu et al., 2016). In general, the total community biomass is observed to increase with both oxygen and methane uptake, as is expected from the increased abundance of nutrients. The community is completely dominated by *Methylobacter* under low methane/low oxygen conditions. An increase of methane at low oxygen conditions does not change the dominance of *Methylobacter*. Furthermore, the increase of oxygen under low methane carbon conditions has minimal impact in changing the community composition, as can be seen from the consistent ratio of *Methylomonas* to *Methylobacter* across the entire range of oxygen uptake at low methane uptake condition (Figure 2). The biggest charge in community composition is observed under high carbon and high oxygen conditions in which the community biomass is composed of 77% *Methylobacter* and 23% *Methylomonas*, as compared to *99% Methylobacter* and 1% *Methylomonas* in the low methane/low oxygen condition (Figure 2). The complete numerical results are included in Supplementary information 4.

In nutrient-rich condition (shown in Figure 3), *Methylobacter* is the dominant community member and consumes the major portion of all of the shared the metabolites *i. e*., methane, oxygen, and nitrogen. *Methylomonas* does not consume a lot of methane and all of it is accumulated in biomass and therefore no formaldehyde is produced. Methylobacter also has a very active serine cycle converting formaldehyde into the metabolites from the central carbon metabolism.

Under the oxygen limiting condition (Figure 4A), the flux through the methane oxidation pathway substantially decreases in *Methylobacter*. However, *Methylobacter* has a higher biomass formation, possibly due to the increased uptake of formaldehyde produced by methane oxidation in *Methylomonas. Methylomonas* consumes the majority (99%) of the total methane uptaken by the community under the oxygen limiting condition (Figure 1B). *Methylomonas* diverts central carbon compounds to produce pyruvate via the assimilation of carbon di-oxide and acetaldehyde under the oxygen limited condition (Figure 4B). A small fraction of carbon di-oxide downstream of this reaction is secreted into the environment, which is the lowest among all the conditions. Additionally, the alpha-ketoglutarate dehydrogenase enzyme becomes active in converting alpha-ketoglutarate into succinyl-CoA (Figure 4B) instead of succinate being produced by succinyldihydrolipoamide in nutrient-rich condition (Figure 3).

The methane oxidizing pathway and TCA cycle reactions in *Methylobacter* has significantly increased fluxes under nitrogen limited conditions (Figure 5A). Specifically, reactions that produce succinate increases by 1.88 mmol/gDCW. *Methylobacter* additionally excretes more carbon di-oxide while consuming more oxygen under nitrogen limited conditions (Figure 1C). On the other hand, the activity of the methane oxidation pathway of *Methylomonas* decreases significantly (the lowest in any non-carbon limiting conditions). Also, succinyl-CoA in *Methylomonas* is produced from alpha-ketoglutarate in the TCA cycle, similar to oxygen limited conditions (Figure 5B).

In the carbon limited growth condition, *Methylomonas* takes up very small amount of methane, while *Methylobacter* takes up most of the methane supplied to the community (Figure 6). The central metabolism of *Methylobacter* is not altered under this carbon limited condition (Figure 6A). On the other hand, in *Methylomonas*, carbon di-oxide was scavenged by assimilating succinyldihydrolipoamide and carbon di-oxide to succinyl-CoA (Figure 6B). This reaction is inactive in the oxygen and nitrogen limiting conditions. *Methylomonas* also displays minimal activity in its serine cycle under every limiting condition.

### Dynamic shifts in metabolism under emulated natural and synthetic community composition

Previous studies have shown inconsistencies between the natural Lake Washington community and synthetic community cultured in the lab. In the natural community, *Methylobacter* has been shown to be dominant under both low methane/high oxygen and high methane/low oxygen conditions (Beck et al., 2013). However, *Methylomonas* outcompeted *Methylobacter* in both low methane/high oxygen and high methane/low oxygen conditions in synthetic co-cultures (Yu et al., 2016). To elucidate the metabolic flux distributions and the extent of inter-species interactions that gives rise to the observed community composition in the natural and synthetic communities, the experimentally observed species abundance ratio was imposed on the growth rates of *Methylobacter* (MB) and *Methylomonas* (ML) as a constraint in the community optimization framework. For naturally occurring community, a MB:ML ratio of 9.3:1.0 and for a synthetic community, a MB:ML ratio of 0.05:1.0 were used.

**Figure 7:** Flux distribution for select metabolites in *Methylobacter* and *Methylomonas* under A) synthetic co-culture conditions and B) natural Lake Washington conditions.

The vast majority of reactions in *Methylomonas* has lower flux values in the natural community compared to the synthetic community. This occurs because *Methylomonas* constitutes a smaller portion of the total community biomass in natural communities. There are alternate pathways to produce pyruvate which has increased flux (Figure 7A). These pathways produce pyruvate by assimilating carbon di-oxide and acetaldehyde and by the assimilation of cysteine and mercaptopyruvate. However, *Methylomonas* produces less pyruvate overall even with the increased flux in these pathways, since other pyruvate producing pathways decrease in flux. Primarily, the flux through L-serine and ammonia assimilation to produce pyruvate is high in the synthetic condition but decreases in natural condition. Under natural conditions, *Methylobacter* uses methane as its primary carbon source while still consuming formaldehyde. On the other hand, *Methylobacter* solely relies upon the secreted formaldehyde from *Methylomonas* as a carbon source under the synthetic condition (Figure 7B).

## Discussion

Aerobic methane oxidation in freshwater lakes around the world is a key metabolic process that significantly affects the carbon cycle by acting as a major sink of methane. Lake Washington provides an wonderful opportunity to study the methane cycling, with up to 20% of the organic carbon being released as methane through decomposition and consuming up to 10% of the dissolved oxygen in the lake water (Kuivila et al., 1988). With the goal to understand the dynamics of the methane-recycling Lake Washington community, we integrated high-quality manually curated and refined genome-scale metabolic models of highly abundant, functionally important, and representative community members using multi-level optimization-based frameworks. While this community have been studied by many researchers in the previous years with *in vivo* tools like synthetic co-cultures, metagenomics, and metatranscriptomics (Hernandez et al., 2015; Oshkin et al., 2015; Yu et al., 2016; Krause et al., 2017), an *in silico* approach like the one used in this work is needed to have a deeper understanding of the underlying mechanisms that govern the inter-species interactions and in turn, the community structure, function and dynamics in Lake Washington. As the global carbon transactions are changing and the release of greenhouse gases in the atmosphere is consistently getting worse every year, it is imperative for us to put our best efforts in mitigating the harmful effects. To do that, the use of genome-scale metabolic modeling tools to understand how the microbial communities are involved in these metabolic processes function in natural environments is essential.

To make the community model a good representation of the naturally occurring methane-recycling community, we have selected two highly abundant and functionally important microbial species. Following the manual curation process of both metabolic models, it was found that *Methylobacter* was more efficient than *Methylomonas* at producing biomass when simulated under the same standard growth environment and biological constraints in the Lake Washington. Since *Methylobacter* and *Methylomonas* are competitors for the sole carbon source, methane, the overall stoichiometries of biomass precursors are important factors in the methane utilization ratio of the two species. While there is no direct literature evidence that suggests that one of them is more efficient in utilizing methane for growth compared to the other, it is highly possible that *Methylomonas* is limited by other small molecules that inhibit high methane consumption, while *Methylobacter* is allowed to take up the major portion of the methane available in the environment. For example, the community is tested under carbon, oxygen, and nitrogen limited conditions to observe how its central metabolic pathway and the community composition varied, and we observe the dominance of *Methylobacter* in the community in all of the growth conditions (Figures 1 through 6). This is observed even during the oxygen limited growth condition, where *Methylobacter* is unable to consume as much methane as *Methylomonas*. In this condition, Methylomonas takes up more methane than Methylobacter, but still in a very low rate. On the other hand, *Methylobacter* can still maintain its dominance, by consuming the formaldehyde produced by *Methylomonas* as a carbon source. Specifically, *Methylobacter* consuming formaldehyde allows it to bypass the oxygen-intensive reaction of oxidizing methane to methanol. This commensal relationship helps *Methylobacter* to enhance its biomass even when the uptake of the original carbon source (methane) is low while protecting *Methylomonas* from the toxicity and growth inhibitory effects of formaldehyde (Hou, Laskin & Patel, 1979; Costa et al., 2001). In our optimization formulation, the inhibitory formaldehyde constraint placed on *Methylobacter* makes the consumption of formaldehyde detrimental towards biomass production but at the same it can act as a carbon source and compensates for its inhibitory effects.

Each of the nutrient limited conditions shows variable differences within the methane oxidation pathway, the serine cycle, and the TCA cycle. While we do not observe a noticeable flux through the serine cycle in *Methylomonas*, it is highly active in *Methylobacter* in all conditions, as it is fed from the formaldehyde either converted from methane (in all conditions except oxygen limiting) or supplied by *Methylomonas* (in oxygen limiting condition). In oxygen limiting condition, the community is also observed to conserve as much resource as possible. For example, *Methylomonas*, while excretes a lot of carbon dioxide in all conditions, routes most of it back to central carbon metabolism through the direction reversal of pyruvate decarboxylase. While there is currently no experimental studies pointing to this phenomenon, a number of studies in other organisms suggest a high oxygen sensitivity of this enzyme (Tadege, Brändle & Kuhlemeier, 1998; Eram & Ma, 2013). Therefore, a possible explanation of the shifts in pyruvate metabolism is the oxygen sensitivity of this enzyme, which needs to be further studied.

Another interesting observation from this study is the activity levels of alpha-ketoglutarate dehydrogenase enzyme in nutrient rich vs. nutrient-limiting conditions. The presence and expression of alpha-ketoglutarate dehydrogenase in type-I methanotrophs like *Methylomonas* has been a matter of debate for quite some time (Zhao & Hanson, 1984; Theisen & Murrell, 2005), which is manifested as the inability of methanotrophs to grow on multi-carbon substrates (Smith, Trotsenko & Murrell, 2010). In this study we observe alpha-ketoglutarate dehydrogenase activity only in oxygen and nitrogen-limiting conditions, while there is no flux through it in carbon limiting condition. At this point it is not straightforward to decipher what exactly might be the regulating factor to this enzyme and warrants further experimentation.

Finally, the community model is simulated with fixed abundance ratios of the two members to reflect the composition in synthetic and natural communities. The changes in community composition as oxygen and carbon levels change are more consistent with behaviors in natural communities than synthetic co-cultures (Hernandez et al., 2015; Yu et al., 2016). *Methylobacter* is able to utilize methane to produce biomass at a more efficient level than *Methylomonas*. However, when the synthetic co-culture condition is imposed on the species abundance ratio, *i.e.*, favoring *Methylomonas* biomass, *Methylobacter* is forced to not take up any methane from the environment and consume only the formaldehyde produced by *Methylomonas*. This is contradictory to what was observed in the natural community and synthetic co-culture experiments (Yu et al., 2016). However, it should be noted that, the synthetic co-culture experiments reported in literature involved other community members, whose metabolic interactions with *Methylobacter* and *Methylomonas* are not well characterized to date. Some studies have indicated co-operative relationships between *Methylomonas* and other species (Beck et al., 2013), which might potentially impact that community dynamics in synthetic co-cultures. Also, despite the dominance of *Methylomonas* in the synthetic community at a species level (Soni et al., 1998; Hoefman et al., 2012), further assessment of community composition at a higher taxonomic level indicated a consistency with naturally observed composition (Yu et al., 2016). We hypothesize that this discrepancy is possible, given that there is high functional redundancy present in Lake Washington community, similar to any naturally occurring microbial ecosystem (Galand et al., 2018; Louca et al., 2018; Islam et al., 2019; Jia & Whalen, 2020).

## Conclusion

In this work, we attempted to enhance our mechanistic understanding of the dynamics in the methane-recycling Lake Washington community through genome-scale metabolic modeling of the representative and functionally important community members, *Methylobacter Tundripaludum 21/22* and *Methylomonas sp LW13.* The understanding of this community behavior will be a foundation for future studies that aim at the long-term goal of creating a complex synthetic community capable of carrying out certain desired functions through the consumption of methane, thus mitigating the harmful effects of methane release in the atmosphere. One should be aware of the fact that the *in silico* results need to be further tested and confirmed through new experiments before any engineering strategies can be successfully employed. The community metabolic model, in that regard, should be expanded to include other major players of the Lake Washington community, *i.e*., members of *Bacteroidetes, Proteobacteria*, and *Betaproteobacteria* phyla. Including these organisms in our community metabolic model will not only enable us to explain currently unidentified inter-species metabolite exchanges/interactions that play important role in the cycling of methane as well as other nutrients but also will be able to provide a realistic representation of the natural Lake Washington community.

